# Genetic diversity and population structure of pedunculate oaks (*Quercus robur*) in Wytham Woods

**DOI:** 10.1101/2024.10.28.619668

**Authors:** Tin Hang Hung, Elias Formaggia, Lucy Morley, Keith Kirby, Roberto Salguero-Gómez, Ben C. Sheldon, John J. MacKay

## Abstract

Genetic diversity is fundamental for adaptation to changing environments. It is particularly important in forest trees because of their significant role in nature’s contribution to people. However, their genetic diversity has been significantly changed by human activities in the past centuries.
This paper investigates a focal site, the Wytham Woods, one of the most researched woodlands on Earth, and presents a population genetic study on pedunculate oaks (*Quercus robur*), a keystone species in the ecosystem. We characterised 210 trees with Genotyping by Sequencing (GbS) and quantified levels of genetic diversity across stands with different histories and management regimes.
We detected only a weak population structure within the 218,567 SNPs, such that most genetic variation occurred within but not among stands, which included semi-natural woodland areas and plantations aged between 200 to 50 years ago. We also observed little difference in observed and expected heterozygosity among stand types, but detected some inbreeding in the youngest plantation. We discovered 26,174 SNPs (11.98%) that were highly differentiated and under potential selection.
We suggest that the life history traits of oak contribute to its resistance against genetic erosion, which is also observed in beeches, spruces, and pines. Preference for oaks as a timber tree and the tendency to use local seed source might have resulted in the homogeneous population structure. However, tree-to-tree differences may harbour variation in putative adaptive loci. Our study contributes crucial baseline information on for conservation and management of human-modified woodlands, in addition to supporting long-term ecological studies on many other species, which depend on this keystone oak species.

**Societal Impact Statement:** Our study highlights the importance of monitoring and preserving genetic diversity in forest trees, particularly in keystone species like pedunculate oaks. Human activities, including land use changes and forestry practices, could influence their genetic diversity and potentially alter nature’s contribution to people. We demonstrate how understanding the genetic structure of oaks in stands ranging from semi-natural to plantations could (1) shed light on the natural history and the consequence of human activities in Wytham Woods, (2) support the many continuing, long-term ecological studies including adaptational potential in the oak population, and (3) be translated for other co-occurring species and other woodlands for effective genetic monitoring at large.

## Introduction

Genetic diversity is fundamental for the adaptation of populations to changing environmental conditions, underpinning species’ adaptability, persistence, and evolution^1,2^. Genetic diversity is particularly important for current populations, as new challenges emerge, such as pests, diseases, climate change, inbreeding, and habitat degredation^3,4^. Understanding genetic diversity in forest trees is of broad importance, as they have a significant role in maintaining biodiversity^5^, supporting community structure^6^, and stabilising ecosystem properties in natural environments^7^, in addition to sustaining productivity and resilience in forestry^8^. Thus, genetic diversity in trees represents an essential aspect of nature’s contribution to people^9^.

Trees are among the most genetically diverse organisms on land^10^. From an evolutionary perspective, they share unique life history traits and characteristics, including high levels of genetic diversity but usually low nucleotide substitution rates^10,11^. Trees are predominantly outcrossing and thus maintain a high level of gene flow through efficient dispersal mechanisms^12^, self-incompatibility^13^, and severe inbreeding depression^14^. Spatial analyses of population genetic structure in long-lived, woody trees reveal that most of species have a higher within-population variation and a lower among-population variation compared to herbaceous and annual plants^15^.

Forest trees are largely undomesticated but, human activities have influenced them with potential impacts on their adaptability. For example, forest management practices have modified tree density and demographics, connectivity, and effective population size leading to effects on their genetic diversity^16^. In Europe, there is a long history of over-exploitation and loss of forest cover during medieval and early modern times^17,18^. Forest cover has increased from the mid-20^th^ century through both artificial and natural regeneration^7^. Simultaneously, human activities, such as species introduction, pollution, and urbanisation, have exerted novel selective pressures with potentially long-term negative effects on fitness^19^.

It has been speculated that forest plantations may have adverse genetic impacts on wild populations and pose genetic risks that are largely neglected^20^. Potential consequences could include loss of genetic variation, breakdown of adaptations, changes to genetic composition within populations, and breakdown of population structure^21^. In England, the lack of genetic diversity in tree planting stocks and its potentially damaging impacts on forest health has led to renewed interest in understanding the genetic basis of resilience in trees^22^. However, a global synthesis on tree species have also shown cases where there is no significant erosion of genetic diversity despite widespread plantations^23^.

### Wytham Woods: A Living Laboratory

Wytham Woods are among the world’s best studied ecosystems and have been part of the UK Environmental Change Network since 1992, with environmental and species population data continuously monitored for over 70 years^24^. Research conducted in Wytham Woods has contributed significantly to understanding wildlife populations and ecosystem dynamics, including the impact of climate variation on badgers^25^, carbon dioxide fluxes in deciduous woodlands^26^, and climate-driven responses in great tits^27^, among many others. One-third of its area is characterised as ancient woodland traditionally managed by coppicing, where pedunculate oaks (*Quercus robur*) are present as standard trees^28^. Alongside with the oaks, the woodland is dominated by European ash (*Fraxinus excelsior*), sycamore maple (*Acer pseudoplatanus*), birch (*Betula* spp.), and European beech (*Fagus sylvatica*)^29^.

Characterising the patterns of genetic diversity in Wytham Woods oaks is important for three main reasons^29^. First, oak has been the main timber tree in the Woods for centuries, but its importance is declining as a consequence of changes in land use and management regimes, which may impact its genetic diversity^30^. Second, mature oak trees in the Woods display spatial variation in crown health^31^ due to a form of chronic oak decline, in which the crown condition is seen to gradually decline over decades^32^. Genetic differences may underpin the variation in susceptibility and resistance to the chronic oak decline^33^. Third, substantial and consistent tree-to-tree variation has been reported in tree phenology^31^, which is likely due at least in part to genetic differences among individuals^34^. Variation in oak phenology has implications for the life cycles in birds and insects, which have been particularly well studied at Wytham Woods^31^. In contrast, the oaks themselves have remained comparatively understudied in this ecosystem.

The overarching aim of this study is to analyse the spatial pattern of genetic diversity across different stand types in *Q. robur* in Wytham Woods, which has adjacent semi-natural woodlands and plantations aged 200 to 50 years ago. We utilise a new reference genome assembled under the Darwin Tree of Life project and analyse 210 individual trees from five stand types across the Woods by using Genotyping by Sequencing (GbS), which has produced more reliable markers than RAD-seq^35^. First, we analyse the population genetic structure. Second, we calculate and compare the genetic diversity among different stand types. Third, we scan for genome-wide outlier signals which may imply selection.

## Methods

### Study site, study species, and sampling

Wytham Woods is a *c*.400 ha woodland in Oxfordshire, United Kingdom (∼51.77° N, 1.32° W). Foliage of *Q. robur* was sampled across 210 trees from a broad range of semi-natural stands and plantations within the woods between June and September of 2023 (**NCBI BioSample records under BioProject PRJNA1135559**). For each tree, 3–5 healthy, non-senescent leaves were collected using either extendable secateurs or an arborist slingshot. Foliage for each sample was placed in a paper envelope, kept in a sealed plastic bag containing silica beads for desiccation and preservation, and then moved to a –20°C freezer at the end of the day.

The oaks sampled are from five different main stand types across the Woods (**Figure 1**): (N) ancient semi-natural woodland where trees are self-sown; (M) mixture of self-sown and planted oaks; (P-E19) early 19^th^-century plantation; (P-M19) mid-to-late 19^th^-century plantation; and (P-M20) mid 20^th^-century plantation.

**Figure 1.**
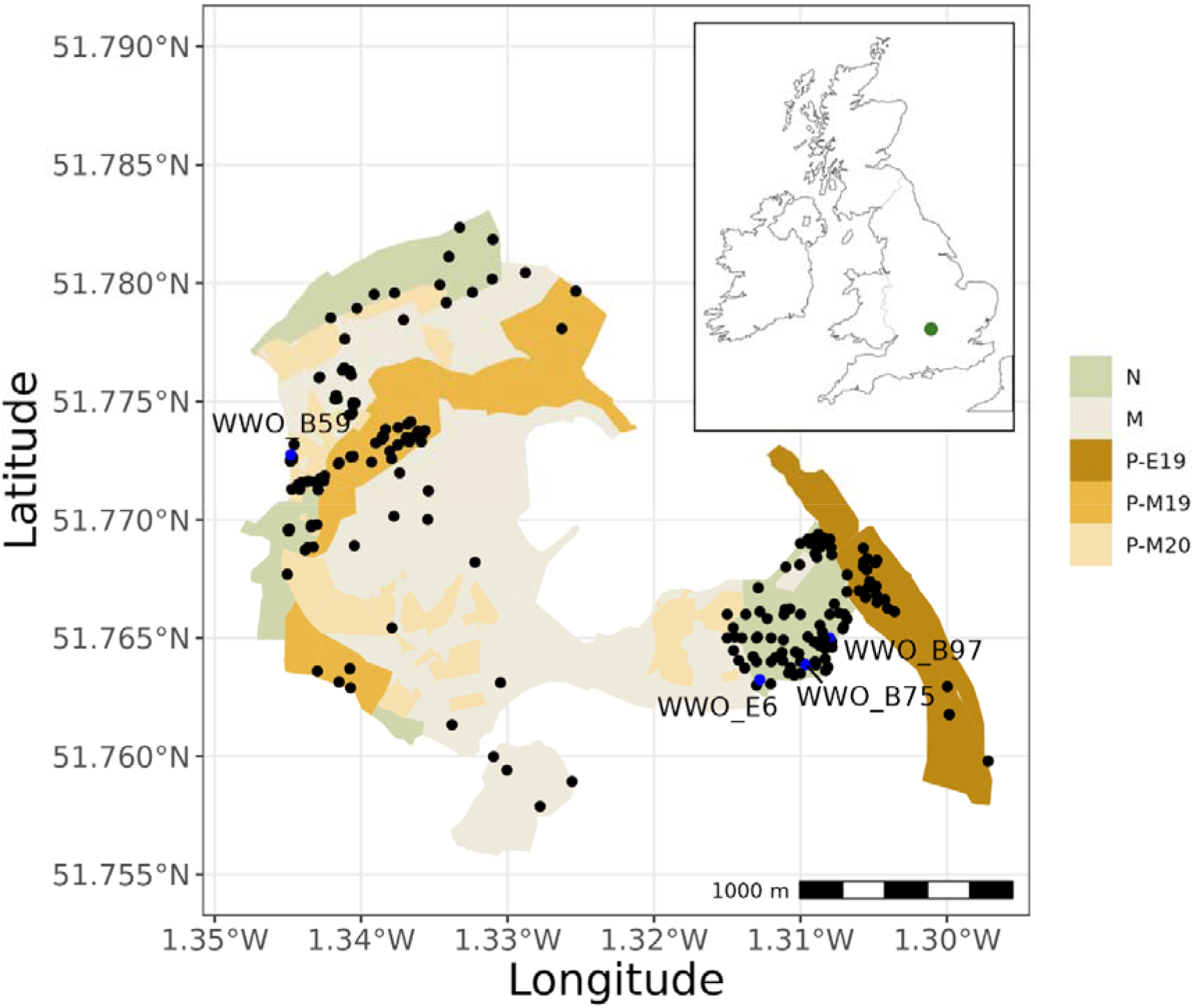
Map of Wytham Woods showing different woodland stand types (N = Semi-natural woodland, M = mixed, P-E19 = early 19^th^-century plantation, P-M19 = mid 19^th^-century plantation, P-M20 = mid 20^th^-century plantation). The inset map at top right corner shows the approximate location of Wytham Woods. Black dots represent individual trees and blue dots represent trees that show level of relatedness implying clonality.

### Library preparation and GbS

Dried foliage tissues were ground using a Retsch TissueLyser II (Qiagen, United Kingdom) at a frequency of 25 s^−1^. Genomic DNA was isolated and purified by using the DNeasy Plant Mini Kit (Qiagen, United Kingdom), according to the protocol with two slight modifications for effective lysis of plant tissues. First, a mix of 370 µl of Lysis Buffer and 30 µ l of protease were used at the lysis step. Second, the incubation time was increased to 1 hour. DNA quantity and quality were assessed using NanoDrop 2000 (Thermo Scientific, United States) and Qubit dsDNA Broad Range assay respectively (Thermo Scientific, United States).

Samples of ∼200 ng genomic DNA were normalised in 10 µL of TE buffer and sent to the Genomic Analysis Platform, Institute of Integrative and Systems Biology, Université Laval (Quebec, Canada) for the preparation of GbS libaries. DNA was digested with PstI (CTGCA↓G), NsiI (ATGCA↓T), and MspI (C↓CGG) according to the 3D-GbS protocol for library preparation^36^. Sub-libraries were barcoded and pooled to equimolarity, then sequenced on an Illumina NovaSeq 6000 system with paired-end mode of 150 bp at the Génome Québec (Montreal, Canada).

### Variant calling

Raw reads were demultiplexed with Sabre 1.0^37^ and trimmed with Cutadapt 1.18^38^ to remove the adaptors and sequences shorter than 50 bp. Reads were aligned against dhQueRobu3.1 (GCA_932294415.1), the reference genome of *Q. robur* from the Darwin Tree of Life project, using bwa-mem 2.2.1^39^. The alignments were indexed and sorted using SAMtools 1.12^40^. Variant pileup and calling were performed with bcftools 1.19^40^. Variants were filtered with proportion of missing data of 0.2 and MAF of 0.01 using VCFtools 0.1.16^41^. Linkage disequilibrium was estimated using PLINK v1.90b6.21^42^, and one SNP per pair was removed based on a *r*^2^ threshold of 0.5 in a genomic window of 50 Kbp and a sliding window of 5 bp.

Estimates of heterozygosity values from SNPs are-context dependent due to sample size and population differentiation^43,44^. Thus, while the filtering above produces sufficient markers for assessing population structure, we also curated another set of SNPs for estimating ‘autosomal heterozygosities’^45^. Variants were filtered so that no missing data were allowed but included all monomorphic markers for unbiased estimation of genetic diversity.

### Estimation of inter-individual relationships

Pairwise relatedness of individual trees was assessed using VCFtools 0.1.16^41^ based on the KING inference (--relatedness2). According to the manual, estimated kinship coefficients of > 0.354, 0.177–0.354, 0.0884–0.177, and 0.0442–0.0884 correspond to duplicate or monozygotic twin, 1st-degree, 2nd-degree, and 3rd-degree relationships, respectively. We also assessed the spatial correlation in the genetic relatedness among individuals by computing Spearman’s rho between pairwise genetic relatedness and geographic distance, where geographic distance was calculated using sf 1.0-17^46^.

### Population genetic structure

We ran sparse non-negative matrix factorisations (sNMF) algorithm in the LEA 3.10 R package^47^ to estimate the number of discrete genetic clusters (*K*) using 10 repetitions for each value of *K* from 1 to 10. This algorithm was considered more statistically robust to departures and violations from classical population genetic model assumptions and computationally efficient than likelihood-based approaches such as STRUCTURE and ADMIXTURE^48^. The optimal *K* was selected for the lowest cross entropy. We also conducted population structure-based outlier analysis with sNMF to detect outlier SNPs that were significantly differentiated among genetic clusters, based on estimated *FST* values from the ancestry coefficients. Multiple testing was corrected with the Benjamini-Hochberg method to obtain *Q-*values for each SNP, which were then mapped across the chromosomes of *Q. robur* and plotted on a Manhattan plot. We also imputed missing genotypes using ancestry and genotype frequency estimates from the sNMF run.

We conducted analysis of molecular variance (AMOVA) to partition the genetic variation within and among stand types using poppr 2.9.6^49^, with 1,000 replications for the Monte Carlo procedure.

### Genetic diversity among different stand types

We calculated the observed heterozygosity (*HO*), the expected heterozygosity (*HE*), and the inbreeding coefficient (*FIS*) using hierfstat 0.5-11^50^ with bootstrapping, for each stand type (N, M, P-E19, P-M19, and P-E20). We also assessed deviations from Hardy-Weinberg Equilibrium (HWE) for each SNP for each stand type using pegas 1.3^51^, with 1,000 replications for the Monte Carlo procedure. We also calculated and reported the autosomal *aHO*, *aHE*, and *aFIS* which consider all polymorphic and monomorphic SNPs for unbiased estimations.

### Annotation and enrichment analysis of highly differentiated genes

We identified SNPs that were both significantly differentiated among clusters in the sNMF test and departed from HWE in each stand type. We used GenomicRanges 1.46.1 to identify their location in the genome and their corresponding gene with the gene annotation of dhQueRobu3.1 (GCA_932294415.1). We also obtained Araport11 annotation^52^ for the gene models in dhQueRobu3.1 with BLASTx 2.11.0+^53^.

We performed enrichment analysis on these highly differentiated genes with clusterProfiler 4.0^54^ on Gene Ontology (GO) terms and Kyoto Encyclopedia of Genes and Genomes (KEGG).

## Results

### GbS results and processing

We obtained 448.10 M × 2 reads in paired-end mode from Illumina GbS, representing 135.33 Gbp in total, from 210 individual oaks. After demultiplexing and cleaning, we retained an average of 2.01 M (± 46.14 K) × 2 paired-end clean reads from each sample, whereas 14.53 M × 2 of sequences were unclassified (3.24%). More than 90% of these reads had quality scores ≥ 30, implying successful data cleaning. The read data were deposited in the NCBI Sequence Read Archive (SRA: SRR29822091–SRR29822300) under BioProject PRJNA1135559. After filtering, we obtained a final set of 218,567 SNPs, which we utilised for pedigree estimations, and analyses of population structure and genetic diversity

### Pedigree estimation

Among the 21,945 (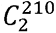) pairwise relationships between the 210 individual trees, most of the pairwise relatedness estimates were negative, with a mean of –0.092 (**Figure 2a**, **Supplementary Table 1**). The vast majority of relationships (98.68%) were classified as unrelated with a > 3°, followed by 3° relationship (0.29%), 2° relationship (0.06%), 1° relationship (0.02%), 0° relationship (0.01%) (**Figure 2b and c**). We detected only two pairs (0.0091%) of clonal trees: WWO_B59 and WWO_E6, which shared a relatedness of 0.43, and also WWO_B75 and WWO_B97, which shared a relatedness of 0.37 (**Figure 1**).

**Figure 2.**
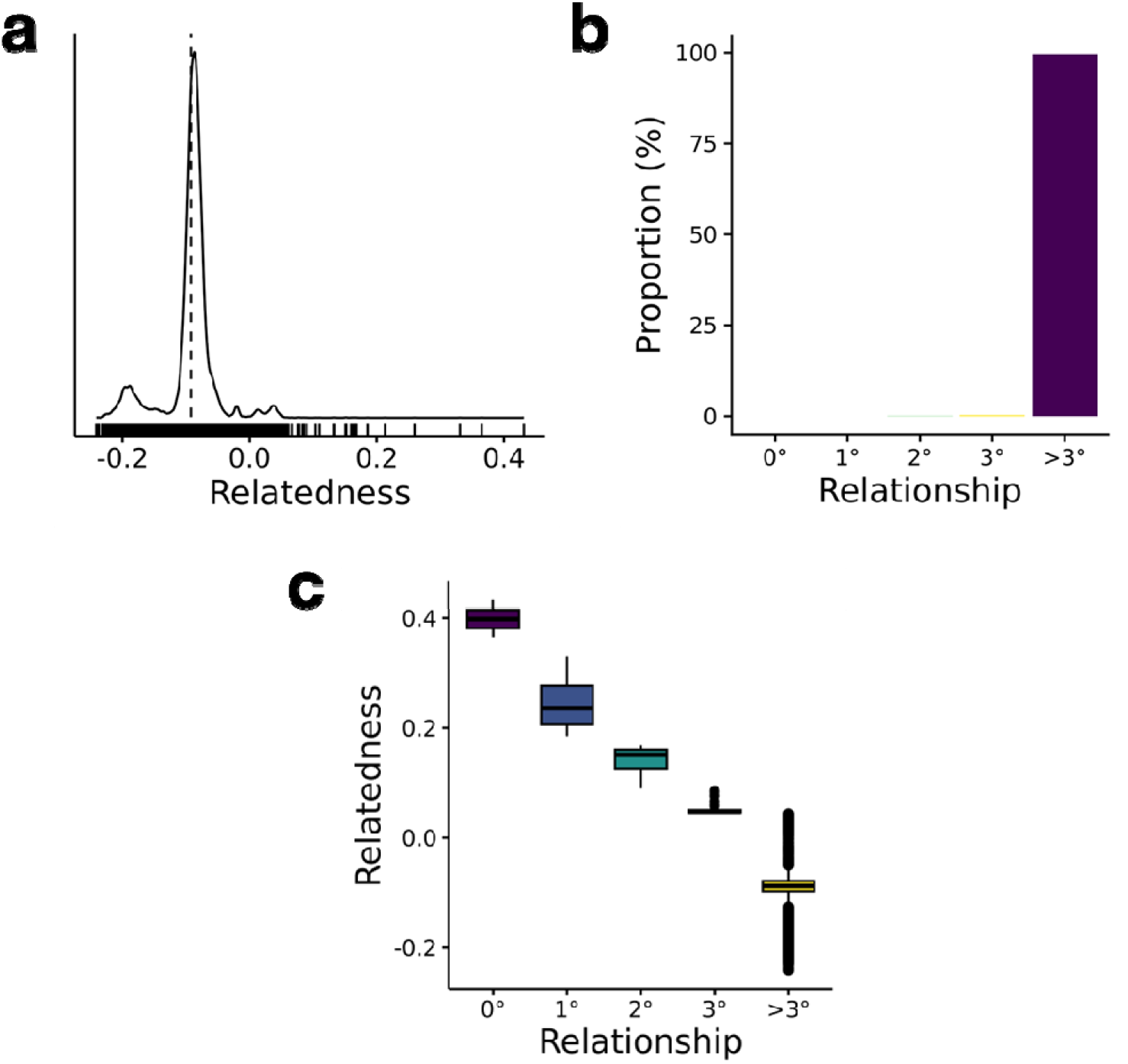
**(a)** Density distribution of pairwise relatedness among individual trees. **(b)** Proportion of pairwise relatedness among different categories of relationships, where relatedness coefficients of > 0.354, 0.177–0.354, 0.0884–0.177, 0.0442– 0.0884, < 0.0442 correspond to duplicate or monozygotic twin (0°), 1st-degree (1°), 2nd-degree (2°), 3rd-degree (3°), and unrelated (>3°) relationships respectively. **(c)** Boxplot of pairwise relatedness among different categories of relationships.

Three trees, 20729, P6B and B199, had a slightly positive average relatedness with all other trees, of 0.0383, 0.0306, 0.0118 respectively (**Figure 3**). All other trees had a negative averaged relatedness with others as expected. There was a very weak negative relationship between the pairwise geographic distance and genetic relatedness (Spearman’s rho = –0.053, *P* = 3.33e–15), suggesting little spatial correlation.

**Figure 3.**
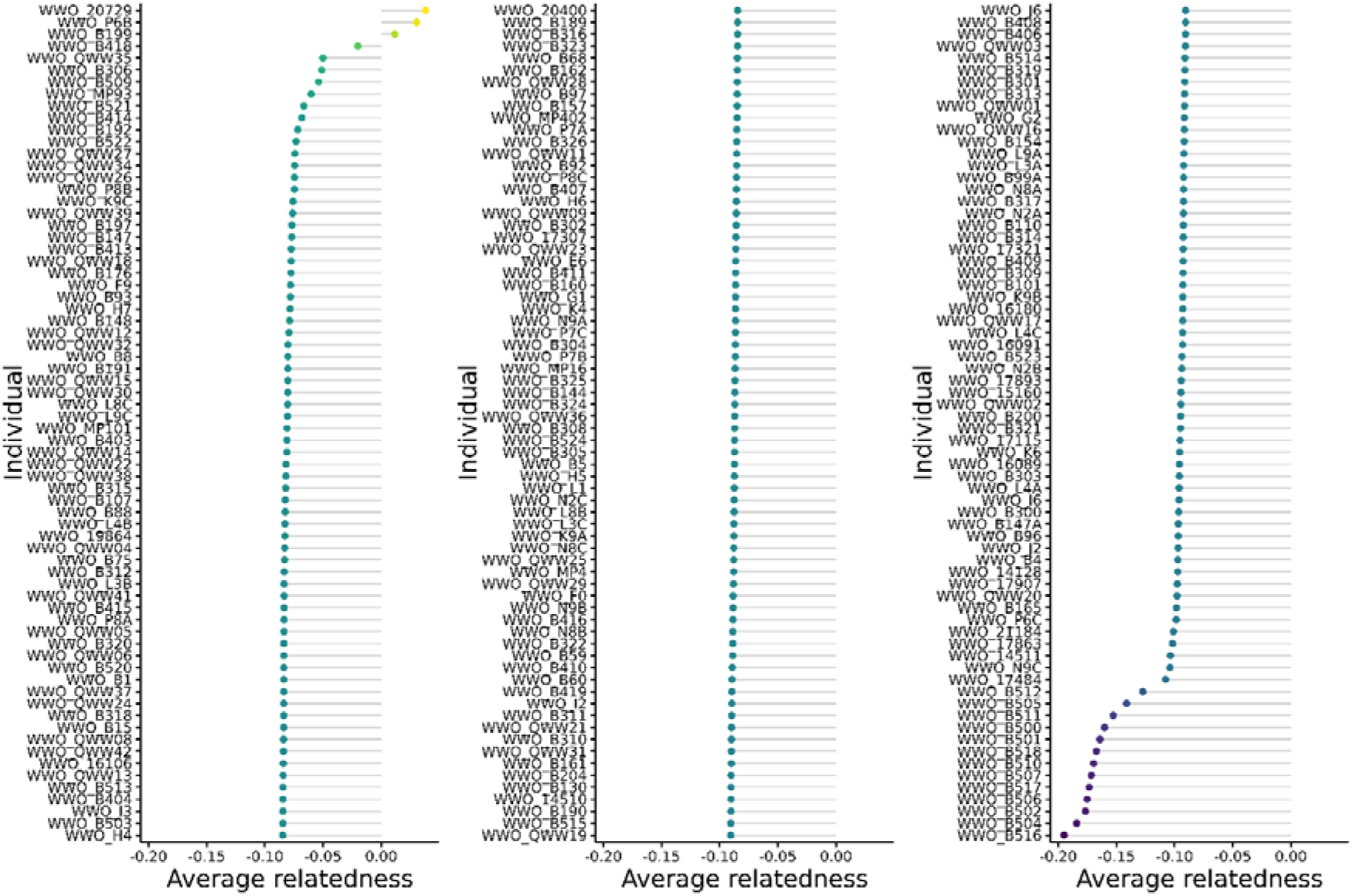
Average relatedness of individual tree with others arranged in descending order, where relatedness coefficients of > 0.354, 0.177–0.354, 0.0884–0.177, 0.0442–0.0884, < 0.0442 correspond to duplicate or monozygotic twin (0°), 1st-degree (1°), 2nd-degree (2°), 3rd-degree (3°), and unrelated (>3°) relationships respectively.

### Population genetic structure

Cross-entropy analysis revealed that *K* = 3 was the most informative clustering in explaining the population genetic structure (**Figure 4a**), while *K* = 2 and 4 were closely similar. The cross-entropy escalated quickly after *K* ≥ 5. However, it appears that there is no clear spatial genetic structure, either with respect to stand types or geographical distance. **Supplementary** Figure 1 visualised the *K =* {2, 3, 4, 5, 6, 7, 8, 9, 10} scenarios. As *K* increased, the genetic clustering broke down for only a small portion of trees, with the remaining large portion unexplained, revealing a very unbalanced hierarchical population structure.

**Figure 4.**
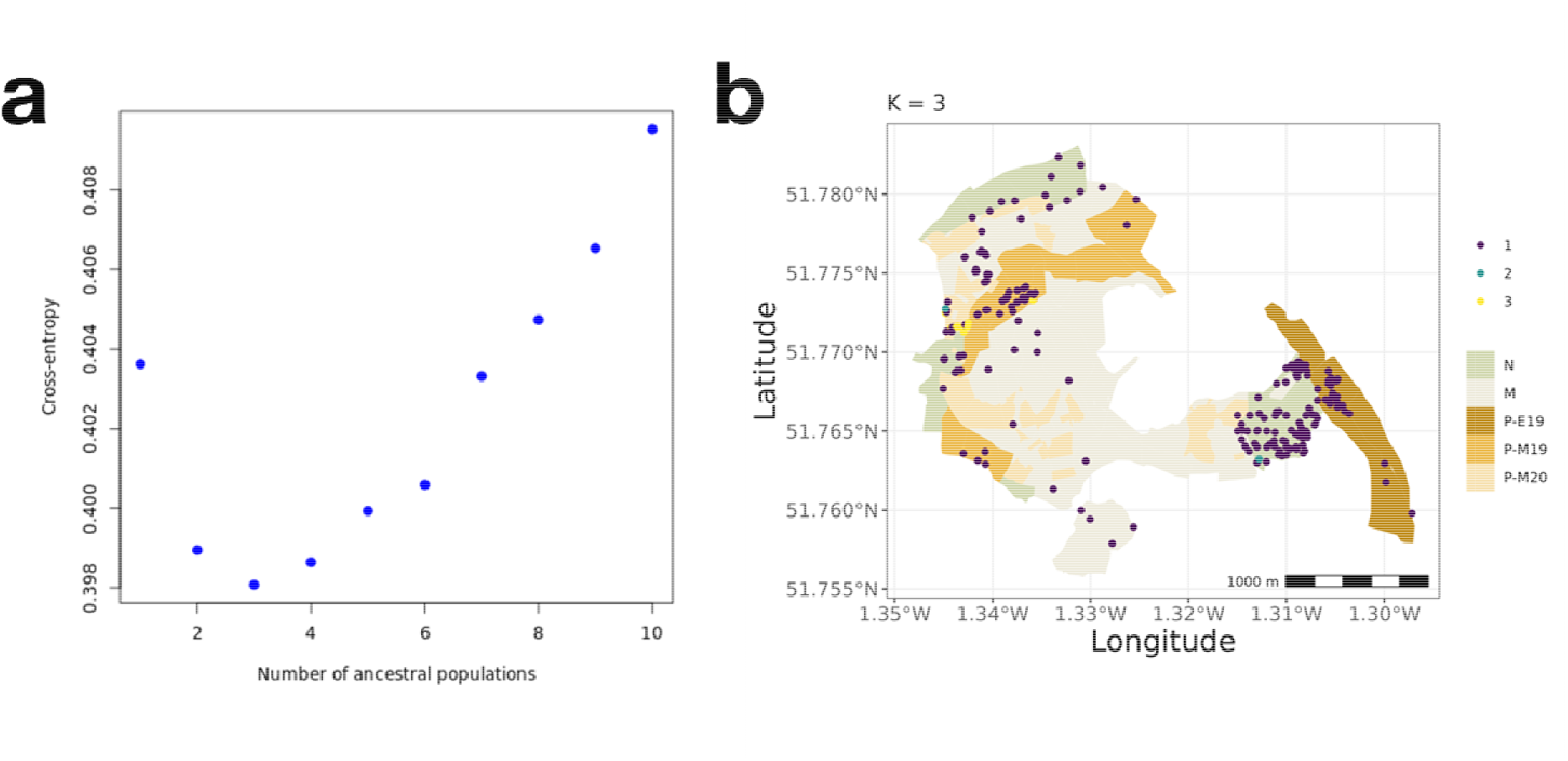
Population genetic structure of pedunculate oaks in Wytham Woods across different stand types and different numbers of clusters *K* predicted by sNMF. **(a)** Cross entropy with increasing *K*. **(b)** The optimal *K* = 3 was determined with the lowest cross-entropy value.

This very weak population genetic structure indicated that there was very little genetic differentiation between the stand types. Only 1.34% of the genetic variation (df = 4, SS = 97481.14, MS = 24370.29) was partitioned between stands, and the other 98.66% (df = 200, SS = 3193578.25, MS = 15967.89) was between trees within stands. The genetic variance components were statistically significant (*P* = 0.001, Monte Carlo test, *n* = 1,000).

### Genetic diversity within different stand types

The levels of observed and expected heterozygosity for oak trees in Wytham Woods were as follows: mean *HO* = 0.119, mean *HE* = 0.133, mean *aHO* = 0.105, and mean *aHE* = 0.102. There was no substantial difference in *HO* or *aHO* among different stand types (**Figure 5a, d**), where M19 plantation had slightly higher values than the others of 0.123 and 0.106, followed by semi-natural woodland (0.121 and 0.105), mixed plantation (0.120 and 0.105), E19 plantation (0.119 and 0.105), and M20 plantation (0.117 and 0.106). There was also no substantial difference in *HE* or *aHE* among different stand types (**Figure 5b, e**), where M19 plantation again had slightly higher values of 0.137 and 0.101, and was followed by M20 plantation (0.136 and 0.108), semi-natural woodland (0.133 and 0.100), mixed plantation (0.132 and 0.101), and E19 plantation (0.130 and 0.101).

**Figure 5.**
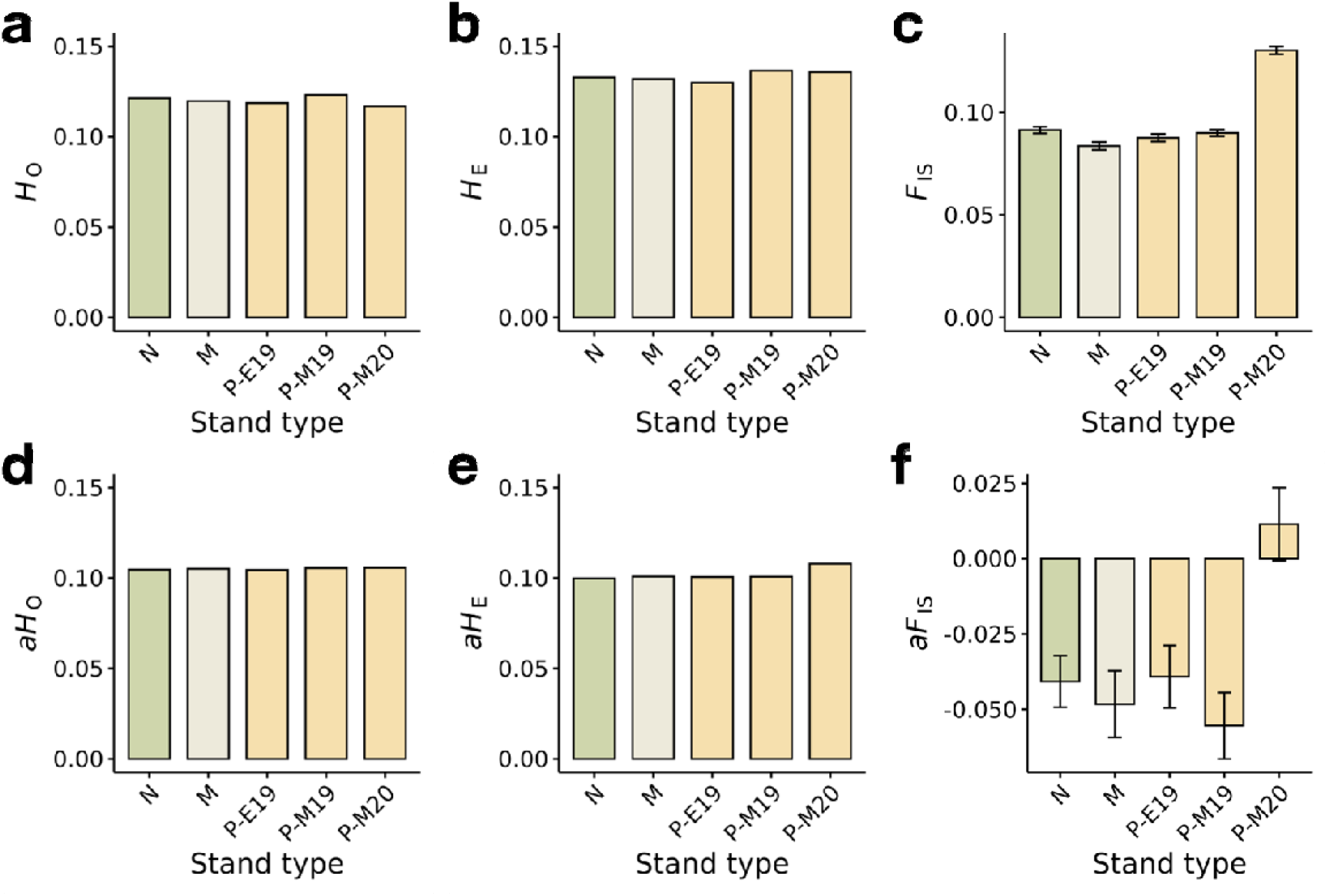
**(a)** Observed heterozygosity (*H_O_*), **(b)** expected heterozygosity (*H_E_*), **(c)** and inbreeding coefficient (*F_IS_*) of all loci and **(d–f)** their autosomal estimates across different stand types: semi-natural (N), mixed (M), early-19^th^-century (P-E19), mid-to-late-19^th^-century (P-M19), and mid-20^th^-century plantations (P-M20). For inbreeding coefficient, the error bar denoted 95% confidence interval determined with bootstrapping.

When using *FIS*, we detected signs of inbreeding in all stand types, where *FIS* differed significantly from zero (95% confidence interval, bootstrapping) (**Figure 5c**). The highest inbreeding was observed in the most recent M20 plantation, with *FIS* of 0.130 (0.128–0.133), followed by semi-natural population (0.0917), M19 plantation (0.0902), E19 plantation (0.0877), and mixed plantation (0.0843). However, when using *aFIS*, we detected no signs of inbreeding as the 95% confidence interval of *aFIS* overlapped with zero in all stand types (**Figure 5f**).

### Identification of putative genes under selection

We detected 40,263 SNPs that were significantly differentiated among the clusters with sNMF algorithm, delineated at the optimal *K* = 3 (*P <* 0.05, after Benjamini-Hochberg correction) (**Figure 6a**, **Supplementary Table 2**). In parallel, we detected 121,219 SNPs that significantly deviated from HWE in at least one of the stand types (*P* < 0.05, Monte Carlo test, *n* = 1,000) (**Supplementary Table 3**). There was a large overlap (26,174 SNPs) among the SNPs sets from the sNMF differentiation and the HWE test results (**Figure 6b**, **Supplementary Table 4**), including 8,287 SNPs with a one-to-one correspondence with 8,287 distinct genes.

**Figure 6.**
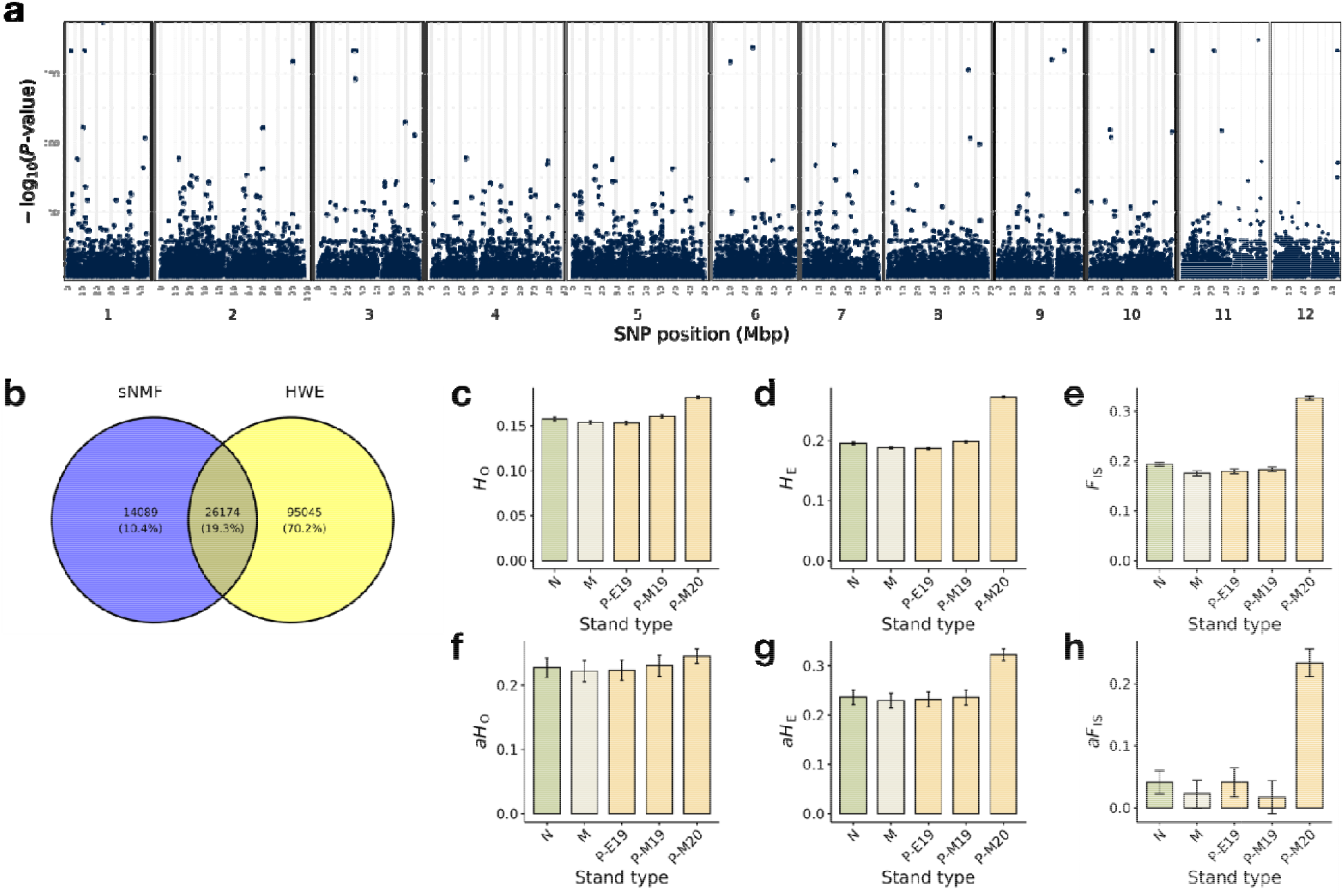
**(a)** Manhattan plot of differentiated loci determined by sNMF. Dark blue dots denote significant loci (*P* < 0.05, after Benjamini-Hochberg correction). **(b)** Venn diagram of differentiated loci determined by sNMF and loci departing from Hardy-Weinberg equilibrium (HWE). (**c**) Observed heterozygosity (*H_O_*), **(d)** expected heterozygosity (*H_E_*), and **(e)** inbreeding coefficient (*FIS*) of loci that were common between sNMF and HWE tests and **(f–h)** autosomal estimates across different stand types: semi-natural (N), mixed (M), early-19^th^-century (P-E19), mid-to-late-19^th^-century (P-M19), and mid- 20^th^-century plantations (P-M20). For inbreeding coefficient, the error bar denoted 95% confidence interval determined with bootstrapping.

The levels of observed and expected heterozygosity for the differentiated loci indicated low to medium levels of genetic diversity but slightly higher than for all loci (mean *HO* = 0.161, *HE* = 0.208, *aHO* = 0.230, *aHE* = 0.251). In addition, there was considerable difference in *HO* and *aHO* among different stand types (**Figure 6c, f**). They were highest in the M20 plantation with 0.182 and 0.245, followed by M19 plantation (0.161 and 0.230), semi-natural woodland (0.158 and 0.228), mixed plantation (0.154 and 0.222), and E19 plantation (0.153 and 0.224). The exact same ranking of values was observed for *HE* and *aHE* (**Figure 6d, g**), where M20 had the highest values and E19 plantation had the lowest.

When considering *FIS* and highly differentiated loci, we detected signs of inbreeding in all stand types, where *FIS* differed significantly from zero (95% confidence interval, bootstrapping) (**Figure 6e**). The highest inbreeding was observed in the most recent M20 plantation, with *FIS* of 0.327 (0.324–0.331), followed by semi-natural woodland (0.195), M19 plantation (0.184), E19 plantation (0.181), and mixed plantation (0.176). However, when considering *aFIS* and highly differentiated loci, we detected signs of inbreeding in all stand types except in the M19 plantation, which had an *aFIS* 95% confidence interval that overlapped with zero. The highest inbreeding was still observed in M20 plantation (0.231), while all others were much lower, only slightly above zero (E19 plantation: 0.042, semi-natural woodland: 0.040, mixed plantation 0.028, and M19 plantation: 0.019)

We were able to assign an annotation to 34,815 out of 35,638 gene models (97.69%) of dhQueRobu3.1 with Araport11. This was followed by enrichment analysis of gene ontology (GO) which identified 183 Biological Process (BP) terms, 29 Cellular Component (CC) terms, and 43 Molecular Function (MF) terms were over-represented among the 26,174 SNPs (*P* < 0.05, after Benjamini-Hochberg correction). Network diagrams showed several interlinked central gene ontology terms. Biological processes included negative regulation of nitrogen compound metabolic process (GO:0010629), negative regulation of gene expression (GO:0051172), gene silencing by RNA (GO:0031047), and DNA repair (GO:0006281) (**Figure 7a**, **Supplementary Table 5**). Cellular components included chromosome (GO:0005694), Golgi apparatus sub-compartment (GO:0098791), trans-Golgi network (GO:0005802), endosome (GO:0005768), and vesicle tethering complex (GO:0099023) (**Figure 7b**, **Supplementary Table 6**). Molecular functions included ATP hydrolysis activity (GO:0016887), ATP binding (GO:0005524), catalytic activity acting on DNA (GO:0140097), ribonucleoside triphosphate phosphatase activity (GO:0017111), and UDP-glycosyltransferase activity (GO:0005524) (**Figure 7c**, **Supplementary Table 7**). In contrast, we detected no significantly over-represented pathway from Kyoto Encyclopaedia of Genes and Genomes (KEGG).

**Figure 7.**
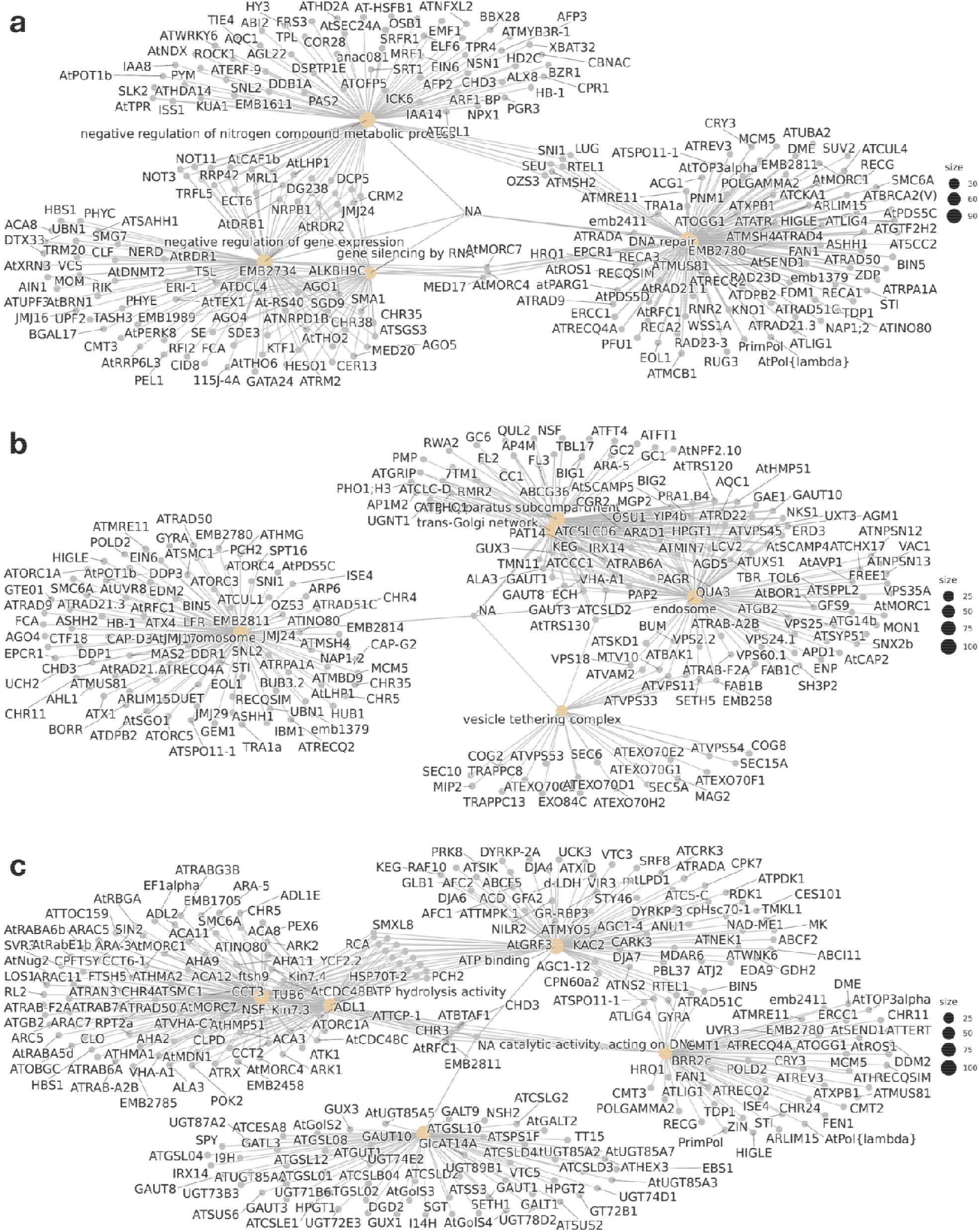
Network plots of over-represented Gene Ontology (GO) terms in categories of **(a)** Biological Processes, **(b)** Cellular Components, and **(c)** Molecular Functions, whereas gene models of dhQueRobu3.1 were annotated with Araport11

## Discussion

Forest trees face a range of challenges from changing environmental conditions and human activities which have been shown to alter their genetic diversity significantly. Understanding of the standing genetic diversity in pedunculate oaks (*Quercus robur*) is thus essential to monitor and predict the adaptability of this species across the shifting climate of the Anthropocene. The present study constitutes the first population genetic study of pedunculate oaks in Wytham Woods, one of the most researched woodlands in the world, and provides a historical perspective of management regimes that may underpin the differences among stand types.

### Small difference in genetic diversity among stand types

We detected minor differences in genetic diversity among the five different stand types of Wytham Woods comprised of semi-natural woodland and plantations of different ages. We only observed a significant signal of inbreeding (*aFIS* > 0) in the most recent plantations established mid 20^th^-century (P-M20). The P-M20 trees possessed an excess of homozygotes (*aHO* << *aHE*) but nonetheless showed the highest *aHO* and *aHE* among all stand types. There are two potential reasons that may explain these observations. First, it may be a classic example of Wahlund effect, where trees (or subpopulations) with different allelic frequencies are combined during the plantation process, thus maintaining a comparable level of genetic diversity as other subpopulations but showing an excess of homozygotes^55^. Second, oak trees only mature and start to produce acorns regularly at 40 to 50 years old^56^. Therefore, the germplasm that was used for establishing P-M20 is likely to come from the youngest trees at the time with limited in reproductive capacity and low levels of gene flow^57^.

It has been widely speculated that forest management practices can have significant effects on genetic diversity and effective population size^23^. However, a global synthesis of research has shown that although genetic diversity can be adversely affected, such as in coppiced oak species (*Quercus* spp.) in the Mediterranean region which have fragmented populations ^58,59^, no decline in genetic diversity was found between semi-natural woodlands and plantations in several cases, such as European beech (*Fagus sylvatica*)^60^, black spruce (*Picea mariana*)^61^, lodgepole pine (*Pinus contorta*)^62^, Douglas fir (*Pseudotsuga menzeisii*)^63^. These species share several important life history traits in common with pedunculate oak including long generation time, longevity, and effective gene flow due to wind pollination, that might have contributed to the resistance in genetic erosion^23^.

Our study also highlighted the importance of standardised methods for the quantification of heterozygosity. At first glance, the genetic diversity of pedunculate oaks in Wytham Woods (mean *HO* = 0.119 and mean *HE* = 0.133) may seem lower than previous reports in the same species, such as in the eastern range of Russia (mean *HE* = 0.669–0.773)^64^, the southern range of Serbia (mean *HE* = 0.740–0.793)^65^ or Croatia (0.195–0.962)^66^. However, such differences are to be expected when contrasting genome-wide SNP marker data as in the present study, and more targeted marker sets, which are usually selected as polymorphic, as reported in oaks including either isozymes^67^, microsatellites (or simple sequence repeats SSR)^64–66^, or small sets (< 100) of SNP markers^68^. The heterozygosity levels cannot be compared directly because of the difference in the marker types. For example, SSRs harbour exceptionally high mutation rates and polymorphism levels when compared to SNPs^69^, and higher estimates of heterozygosity across several species^70–72^.

There are a few tree species that have been assessed with both genome-wide SNP analyses and SSRs for genetic diversity. A GbS study on Norway spruce reports a *HO* of 0.237 and a *HE* of 0.271^73^ for SNPs but the same authors report a mean *HO* of 0.585 and a mean *HE* of 0.722 when using SSR^74^. Similarly, work in Siamese rosewood reported a mean *HO* of 0.56 and a mean *HE* of 0.57 with SSR, but a mean *HO* of 0.18 and a mean *HE* of 0.18 using RAD-seq SNP^75^. Even when considering 130 SNP loci in Russian populations of mixed *Q. robur* and *Q. petraea*, *HO* was between 0.259 and 0.291 and *HE* was between 0.219 and 0.272^76^, providing a strong contrast with SSR findings in the same geographic area. We therefore advise future oak studies to use unbiased parameters, such as autosomal heterozygosities^45^, which consider both monomorphic and polymorphic sites, so that we can continue to monitor and compare genetic diversity meaningfully, especially when they are important parameters in forest conservation and management^77^.

### Weak population structure and history of Wytham Woods

The nature and intensity of woodland management has changed considerably in Wytham Woods, over the last few hundred years as the coppice management of the 18^th^ to early 19^th^ centuries was gradually replaced by high forest and conifer plantations in the 20^th^ century^78^. The oak in the coppice management era was mainly as standard trees, a few of which might be harvested at each coppice cut. There were also occasional much larger fells of oak as indicated in the adverts for local timber sales, but prior to the 19^th^ century there was probably little planting of oak. The fifth Earl of Abingdon, who owned the Woods between 1799 and 1854, then initiated a massive programme of restructuring and new planting under the Enclosure Acts in 1814 and 1816 with some further planting during the later part of the 19^th^ century^78^.

In the 20^th^ century there was heavy felling during the World Wars. After WWII in 1942, the Woods were passed to the University of Oxford. The University’s Department of Forestry organised widespread planting of oak and other species during the 1950s to fill gaps by wartime felling. Since 1960 there has been only small amounts of oak plantings^79^.

History of the woods may explain the weak population structure in Wytham oaks, such that most of genetic variation occurs between individual trees but not among stand types. First, the lack of difference in genetic diversity across stands dating from different periods may reflect a tendency to use local seed sources. Estates often had their own nurseries and might collect acorns from their best stands^80^. Second, despite the overwhelming preference for oak as a timber tree, there may only be strong impact at loci controlling economic traits, maintaining the overall homogeneity of genetic makeup^81^.

### Potential selection pressure on highly differentiated loci

Loci that are highly differentiated among genetic clusters and depart from Hardy-Weinberg equilibrium can offer important insights into natural selection and local adaptation. Given the weak population structure we observed, background levels of *FST* are weak and traditional outlier test based on the empirical distribution of *FST* might fail to detect loci under selection^82^. Therefore, we utilised an approach that extends the definition of *FST* as the proportion of genetic diversity due to differences in allele frequency among populations in a model with admixed individuals^83^, which are frequently introduced by seedling plantations or direct seeding^84^. Loci that depart from Hardy-Weinberg equilibrium can indicate non-random mating, population structure, and small population^85,86^, however the Wahlund effect from subpopulation structure can be ruled out in the case of Wytham oaks. Thus, it is likely that these common loci between the outlier test and HWE test represent signatures of selection.

Analysing loci under potential selection reveals many overrepresented Gene Ontology (GO) terms, which provide indications for understanding the molecular mechanisms and biological processes that may drive adaptation in Wytham Woods. Enrichment of terms such as “negative regulation of nitrogen compound metabolic process” and “negative regulation of gene expression” highlights the importance of homeostasis and transcriptional regulation. Nitrogen metabolism is crucial for plant growth and development, such as amino acid synthesis and nucleotide metabolism^87^. Regulatory mechanisms on nitrogen metabolism are essential for maintaining metabolic balance and optimise resource use under variable environmental conditions^88,89^. Similarly, regulation is needed to ensure optimal gene expression in adaptive responses of plants to environmental changes^90^.

Overrepresented genes in ribonucleoside triphosphate activity similarly imply that loci involved in nucleotide metabolism and synthesis are under selection. They mediate the synthesis of RNA through transcription^91^. They are also involved in signal transduction, recycling of nitrogen, and modification of oxidative stress^92^, as plants must face a variety of environmental stresses. On the other hand, UDP-glycosyltransferase activity underscores the importance of glycosylation processes in adaptation. UDP-glycosyltransferases regulate the modification of various substrates, including hormones, secondary metabolites, and xenobiotics, thereby altering their solubility, stability, and activity. Modification of secondary metabolites are essential for adapting the physiological processes to the changing environment.

### Implications for conservation and future studies of Wytham Woods

Intraspecific variation, as a major component of biodiversity, has received relatively little attention when formulating conservation plans in wild species and ecosystem services management^93,94^. However, the genetic consequences of land use change and human-induced climate change in many species and population still remains unknown^23,95^. The landmark Kunming–Montreal Global Biodiversity Framework, from Convention on Biological Diversity meeting in 2022 (COP15), mandates countries to monitor, manage, and report on the genetic status of species in local, regional, and global programmes, where intraspecific genetic variation is a key indicator^96,97^.

By combining studies of the genetic diversity in oaks in Wytham Woods with its known history, we are able to suggest possible mechanisms for how changes in land use and forestry practices affect genetic diversity. In return, the many continuing, long-term ecological studies in the Woods may benefit from a better understanding of the genetic variation in a keystone species. Moreover, we believe that our study in genetic diversity of oak trees in Wytham Woods may be relevant for other co-occurring species, opening new research avenues that can benefit both conservation and productive forestry.

## Data availability statement

The read data from Illumina sequencing were deposited in the NCBI Sequence Read Archive (SRA: SRR29822091–SRR29822300) under BioProject PRJNA1135559, along with associated BioSample records.

## Author contributions

T.H.H.: designed the study, conducted the bioinformatic analyses, and drafted the manuscript;

E.F.: collected and processed the samples and revised the manuscript;

L.M.: collected and processed the samples and revised the manuscript;

K.K.: processed the spatial data and revised the manuscript;

R.S.: designed the study, collected the samples and revised the manuscript;

B.C.S.: designed the study, collected the samples and revised the manuscript;

J.J.M.: designed the study, collected the samples and revised the manuscript.

## Acknowledgements

This work was supported by funding to T.H.H., R.S., B.C.S., and J.J.M. from the Oxford University Press John Fell Fund (0013514). We also thank Sienna Rattigan for assistance in the fieldwork.

